# Distinct mechanisms of visual and sound adaptation in the cat visual cortex

**DOI:** 10.1101/2024.04.11.589131

**Authors:** Yassine Yahia Belkacemi, Solène Hospital, Nayan Chanauria, Oliver Flouty, Stéphane Molotchnikoff, Vishal Bharmauria

## Abstract

Sensory areas exhibit modular selectivity to stimuli, but they can also respond to features outside of their basic modality. Several studies have shown cross-modal plastic modifications between visual and auditory cortices; however, the exact mechanisms of these modifications are yet not completely known. To this aim, we investigated the effect of 12 minutes of visual *vs.* sound adaptation [forceful application of a non-optimal stimulus to a neuron(s) under observation] on the infra- and supra-granular primary visual neurons (V1) of the cat (*Felis catus*). Previous reports showed that both protocols induced orientation tuning shifts, but sound increased the bandwidths. Here, we compared visual *vs.* sound adaptation effects, specifically analysing the raw tuning curves by computing the area under the curve (AUC) on a trial-by-trial basis. We report that sound adaptation elicited broader tuning curves accompanied with increased variance in the supra- and infra-granular layers, compared with visual adaptation. These findings suggest unique modulation of dendritic structure by distinct adaptation protocols, resulting in disparate tunings. We suggest that broader tuning curves after sound adaptation may keep the visual cortex prepared across a spectrum of abstract representations that match with visual stimuli.

## 1. Introduction

Sensory areas across species are selective to specific properties of trigger features (Hubel and Wiesel, 1959, 1965; Vidyasagar et al., 1996; Swindale, 1998; Chklovskii and Koulakov, 2004; Tolias et al., 2005). For example, in the visual cortex, neurons are selective to orientation / direction, spatial and temporal frequencies, etc., (Hubel and Wiesel, 1962; Maldonado et al., 1997; Kaschube et al., 2010). In the auditory cortex, neurons are selective to a narrow range of auditory frequencies (Aertsen and Johannesma, 1981; Calhoun and Schreiner, 1998; Brewer and Barton, 2016). Similarly, neurons in another sensory areas also have a preferred stimulus (Taube et al., 1990; Bridgeman et al., 1997; Kitchin, 2001; Andersen and Buneo, 2002; Basole et al., 2003; Eichenbaum, 2015; Bednar and Wilson, 2016; Stachenfeld et al., 2017). Nonetheless, neurons in these areas can also respond to trigger features outside of their basic modality i.e., they exhibit cross- or multi-modality (Budinger et al., 2006; Kayser et al., 2008; Teichert and Bolz, 2018; Deneux et al., 2019; Dall’Orso et al., 2020; Garner and Keller, 2022). For example, neurons in visual areas can respond to auditory stimuli (Ibrahim et al., 2016; Chanauria et al., 2019; Bimbard et al., 2023) and vice versa (Kayser et al., 2008). The activity in the visual cortex is attributed to direct projections from the auditory cortex (Iurilli et al., 2012; Ibrahim et al., 2016; Deneux et al., 2019).

Sensory areas are highly plastic, and several investigations have shown that it is possible to change neuronal properties well into adulthood, using specific experimental protocols such as visual deprivation, environmental enrichment and adaptation (Kohn, 2007; Bharmauria et al., 2022; Tring et al., 2023). Adaptation refers to an imposition of a non-preferred stimulus (adapter) to a neuron (s) under observation for a specific duration of time (Kohn, 2007; Bao et al., 2013; Bharmauria et al., 2022). For example, adaptation in the visual cortex leads to two types of orientation tuning shifts in neurons: attractive and repulsive. Attractive shift corresponds to the shift toward the adapter, where repulsive shift represents the shift away from the adapter. Some neurons may not change their selectivity, referred to as refractory neurons (Dragoi et al., 2000; Ghisovan et al., 2009; Bachatene et al., 2013). Similarly tuning shifts have been reported in the auditory cortex (Gourvitch and Eggermont, 2008; Briley and Krumbholz, 2013; Lanting et al., 2013), the middle temporal (MT) area (Kohn and Movshon, 2003, 2004), and other brain regions (Merzenich et al., 1984; Armstrong-James et al., 1994; Webster and Mollon, 1995; Karni and Bertini, 1997; Harris et al., 2000; Reisenman et al., 2003; Gilbert et al., 2009; Marshansky et al., 2011).

Recently, it has been shown that sound can influence the properties of visual neurons (Iurilli et al., 2012; Ibrahim et al., 2016; Chanauria et al., 2019; Bimbard et al., 2023). In fact, auditory stimuli may potentially offer global inhibition to the primary visual cortex (V1) and convey specific tone-related information, especially when reinforced through extended exposure or specific training (Iurilli et al., 2012; Ibrahim et al., 2016; Meijer et al., 2017; Deneux et al., 2019; Knöpfel et al., 2019; Garner and Keller, 2022; Bimbard et al., 2023). In our experiments, we specifically showed that a mere 12-min presentation of an auditory stimulus comprising several frequencies leads to shifts (on either side of the original peak of the tuning) of tuning of visual neurons (Chanauria et al., 2019). Moreover, in the same investigation, using orientation selectivity index (OSI), we reported that sound adaptation significantly increased the bandwidth of layer 5/6 neurons. On the other hand, for layer 2/3, although the bandwidth decreased but no significance was observed. In fact, such 12-min visual adaptation protocols led to both attractive and repulsive shifts in the V1 of cats and mice without changing the bandwidth and OSI of neurons pre- and post-adaptation (Ghisovan et al., 2009; Jeyabalaratnam et al., 2013; Bachatene et al., 2015c; Chanauria et al., 2016; Ouelhazi et al., 2019). Thus, we hypothesized that auditory *vs.* visual stimuli may have unique influence on the excitatory / inhibitory influence of visual neurons, thereby specifically impinging upon shifts and tuning of visual neurons. This led to the following question: are the mechanisms of adaptation in the visual cortex same / similar for visual *vs.* sound adaptation protocols?

To this goal, we compared visual neurons to 12-min visual *vs.* 12-min sound adaptation. In both cases, neurons exhibited shifts of orientation tuning curves (Ghisovan et al., 2009; Bachatene et al., 2015c; Chanauria et al., 2019) Here, by specifically analysing the raw and centered tuning curves, we demonstrate that after sound adaptation neurons showed on average larger response variance and bandwidths accompanied with higher firing rates, suggesting that the underlying dendritic structure was differentially triggered and recruited in both cases. This broadening of the tuning curves may prepare the brain for a larger repertoire of abstract representations of sound that can be matched with the responses of the visual cortex.

## 2. Methods

### 2.1 Ethical Approval

Thirteen (5 for visual adaptation and 8 for sound adaptation) mature domestic cats (*Felis catus*) of either gender were used for experiments in compliance with the regulations sanctioned by the National Institutes of Health (NIH) in the United States and the Canadian Council on Animal Care (CCAC). The cats were procured from the Division of Animal Resources of the University of Montreal. Surgical and electrophysiological recording procedures on the animals were conducted in accordance with the protocols of Canadian Council on Animal Care protocols and were authorized by the Institutional Animal Care and Use Committee of the University of Montreal (CDEA).

### 2.2 Anesthesia

The cats were initially sedated using a combination of Acepromazine (Atravet, 0.1Lmg/kg, administered subcutaneously, Wyeth-Ayerst, Guelph, ON, Canada) and atropine sulphate (Isopto Atropine, 0.04 mg/kg, administered subcutaneously, Atrosa; Rafter, Calgary, AB, Canada), followed by an anesthetic dose of ketamine (Narketan, 40Lmg/kg, administered intramuscularly; Vetoquinol, QC, Canada). Anesthesia was maintained during the surgical procedure through isoflurane inhalation (2%, AErrane; Baxter, Toronto, ON, Canada). Post-surgery, the cats were secured on the stereotaxic apparatus and rendered immobile by the infusion of gallamine triethiodide (Flaxedil, 40Lmg/kg, administered intravenously; Sigma Chemical, St Louis, MO, USA). Artificial ventilation was upheld using a blend of O_2_/N_2_O (30:70) and isoflurane (0.5 %). The paralysis was sustained by continuously infusing gallamine triethiodide (10 mg/kg/h) in a 5 % dextrose lactated Ringer’s solution (administered intravenously, Baxter, Mississauga, ON, Canada) throughout the duration of the experiment.

### 2.3 Surgery

Xylocaine, a local anesthetic (2%; AstraZeneca, Mississauga, ON, Canada), was injected beneath the skin before any incision during the surgery. A heated pad was positioned under the cat to sustain a body temperature of 37.5 °C. To prevent bacterial infection, antibiotics, Trimethoprim and Sulfadiazine (Tribrissen; 30 mg/kg/day, administered subcutaneously; Schering Plow, Pointe-Claire, QC, Canada), and Benzylpenicillin procaine and Benzylpenicillin benzathine suspension (Duplocillin; 0.1LmL/kg, administered intramuscularly; Intervet, Withby, ON, Canada), were given to the animals. Initially, a vein in the forelimb of the animal was cannulated, followed by a tracheotomy for artificial ventilation. Continuous monitoring of the EEG, electrocardiogram, and the animal’s expired CO_2_:O_2_ saturation ensured adequate anesthesia depth throughout the experiment, while maintaining end-tidal CO_2_ partial pressure between 25 and 30LmmHg. Subsequently, a craniotomy (1*1Lcm square) was conducted over V1 (areas 17/18, Horsley-Clarke coordinates P0–P6; L0–L6). The underlying dura was removed, and a multichannel electrode (several contacts in depth) was placed in the area 17 **(Fig. 1A)**. The pupils were dilated using atropine sulfate (1%; Isopto-Atropine, Alcon, Mississauga, ON, Canada), and the nictitating membranes were retracted using phenylephrine hydrochloride (2.5 %; Mydfrin, Alcon). To prevent corneal dehydration, plano contact lenses with artificial pupils (5 mm diameter) were applied to the cat’s eyes. Finally, at the conclusion of the experiment, the cats were euthanized with a lethal dose of pentobarbital sodium (100 mg/kg; Somnotol, MTC Pharmaceuticals, Cambridge, ON, Canada) administered via intravenous injection.

**Figure 1:**
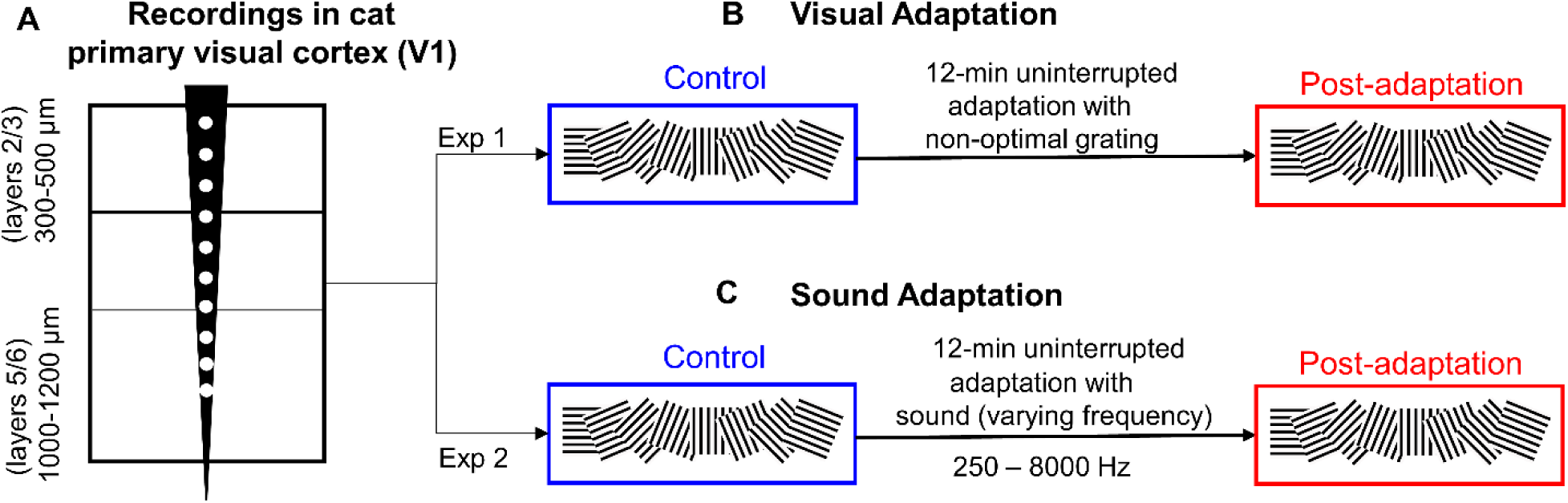
Adaptation protocol for the visual vs. sound adaptation in the cat visual cortex. **(A)** Neural recordings were performed using a depth multielectrode with contacts spanning the supra-((300-500 µm) and infra-granular layer (1000-1220 µm) of the cat V1. **(B)** Exp 1: visual adaptation, multiunit recordings were performed in relation to different (randomly presented) drifting sine wave gratings in the control condition, followed by 12-min uninterrupted adaptation with a non-optimal grating. After adaptation, neural recordings were again conducted in relation to the base set of the oriented gratings. **(C)** Exp 2: sound adaptation, same as B, except the adaptation was done with a repeating sound with varying frequency components from 250-8000 Hz (Chanauria et al., 2019). Note: both protocols only varied in the presentation of adapter.

### 2.4 Stimuli and Experimental Design

Two types of stimuli, visual **(Fig. 1B)** and audio **(Fig. 1C),** were utilized. The visual stimuli were within a 15 ° radius from the fovea, with the receptive fields (RF) centrally located. Monocular stimulation was performed, and the edges of the RF were examined using a handheld ophthalmoscope once clear detectable activity was observed (Barlow et al., 1967). This involved moving a light bar from the periphery toward the center until a response was triggered. The contrast was set at 80 %, and the mean luminance at 40 cd/m^2^. Optimal spatial and temporal frequencies were established at 0.24 cycles/deg. and within the range of 1.0–2.0 Hz, as these values elicit maximal response from V1 neurons when employing drifting sine-wave gratings(Bardy et al., 2006). Subsequently, oriented gratings were randomly presented covering the excitatory RF **(Fig. 1B-C),** to compute the neurons’ orientation tuning curves pre- and post-adaptation in both protocols (Maffei et al., 1973). The visual stimuli were generated using a VSG 2/5 graphics board (Cambridge Research Systems, Rochester, England) displayed on a 21-inch Monitor (Sony GDM-F520 Trinitron, Tokyo, Japan), positioned 57 cm from the cat’s eyes, with 1024 × 768 pixels and running at 100 Hz frame refresh.

#### 2.4.1 Visual adaptation

Figure 1B displays the visual adaptation protocol. The drifting sine-wave gratings were presented unidirectionally in eight possible orientations (base set) and were randomly presented 25 times for 4 s each, with an inter-stimulus interval of 1–3 s. After that a non-optimal orientation was presented for 12 minutes followed by the presentation of the base set again (beginning with the presentation of the adapter). This protocol has been frequently employed in other investigations (Marshansky et al., 2011; Bachatene et al., 2012b, 2015b, 2015a; Chanauria et al., 2016; Ouelhazi et al., 2024).

#### 2.4.2 Sound adaptation

The sound adaptation consisted of different experiments and its effect on tuning shifts has been previously reported (Chanauria et al., 2019). Following the presentation of a base set (Fig. 1C) as for visual experiment, the animal was exposed to broadband noise-like auditory stimuli (adapter) at 78LdB SPL, consisting of temporally orthogonal rippled combinations (TORCs) with varying frequency components from 250L-8000LHz (Chanauria et al., 2019). This 3 s auditory stimulus was played continuously for 12Lminutes using a pair of external loudspeakers positioned at 57 cm perpendicular to the animal’s axis (Fig. 1C). The speakers’ frequency response range was 120 Hz–18 KHz. In some recordings, the speakers were moved laterally at 30 cm on either side from the fixation axis of the cat, aiming to assess response changes with speaker position alterations. Sound frequency and intensity were optimized using Bruel and Kjaer Spectris Group Sonometer according to the experimental design and set on a standard C-scale of the sonometer for both ears. Immediately after the 12-min acoustic stimulus, the orientation base set was again randomly presented. Notably, during sound application, no visual stimuli were presented.

### 2.5 Electrophysiological recordings

The study involved examining multiunit activity within the V1 of anesthetized cats. Neural signals were recorded using a tungsten multichannel electrode from Alpha Omega Co. USA Inc., featuring an impedance range of 0.1 to 0.8LMΩ and a sampling rate of 25,000 KHz. Neural activity was recorded sequentially from both cerebral hemispheres of the cats, exposing one hemisphere at a time. The electrode setup comprised four microelectrodes housed in stainless-steel tubing, arranged linearly with a 500 μm spacing between electrodes (Fig. 1A). Signal acquisition from these microelectrodes was done using Spike2 software developed by CED in Cambridge, UK. The acquired signal underwent amplification and band-pass filtration within the range of 300Hz-3 KHz before digitization. The digitized signal was then displayed on an oscilloscope and recorded with a temporal resolution of 0.05Lms.

In both cases, the recordings were performed at cortical depths averaging between 300–500 μm (supragranular) and 1000–1200 μm (infragranular) simultaneously from both recording sites (Chanauria et al., 2016, 2019). Post-recording, spike sorting was conducted using the Spike2 package from CED, Cambridge, UK. To ensure clean spike sorting, precautions were taken due to potential variations in neuronal waveforms resulting from various factors (Quiroga et al., 2004; Quiroga, 2012). Single units were differentiated using criteria including spike waveforms, principal component analysis (PCA), and autocorrelograms (ACG), as previously detailed (Bharmauria et al., 2016a, 2016b).

### 2.6 Data analysis: area under the curve (AUC) computation

We computed the orientation tuning curves of neurons that have been previously published (Chanauria et al., 2016, 2019). Here, with the specific question in mind, we focused on measuring the variance and tuning of neurons pre- and post-adaptation. We compared the above datasets in three ways: 1) the area of the raw responses (tuning curve) was computed for a neuron on a trial-by-trial basis, i.e., no centering of the response was done. 2) The mean of the 25 trials was computed and then the orientation with the highest mean responses was centered. Then the area under the tuning curve was computed on a trial-to-trial basis. 3) The area was computed on a trial-by-trial basis, where every trial with the maximal response was centered for each neuron.

The area under the curve was computed for each trial using numerical integration via the trapezoidal method using the matlab function (trapz).

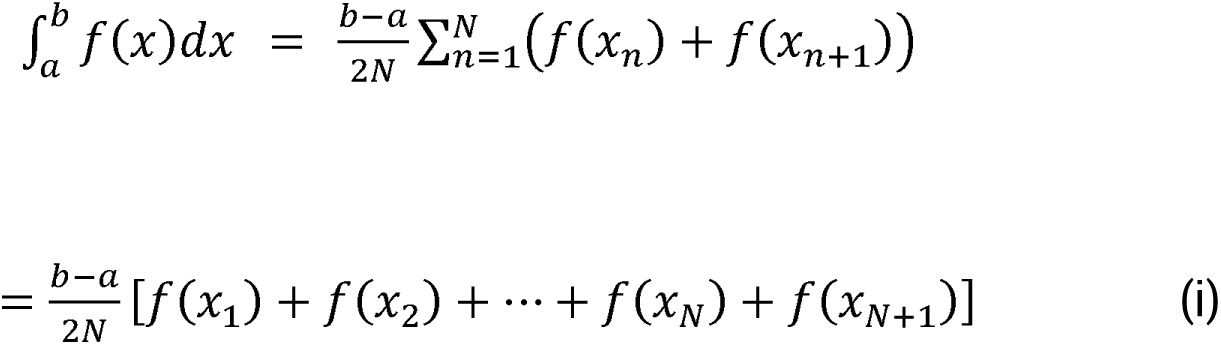

where the spacing between each point is equal to the scalar value 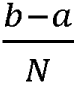. *N* represents the data length which is the number of orientations, *a* represents the first orientation and *b* represents the last orientation. *F(x_n_)* represents the firing rate (counts/bin). The AUC was computed for the raw and centered data for neurons in each layer for the visual and sound conditions. An example of the computation of the AUC is shown in **the Supplementary** Figure 1.

## 3. Results

Recent investigations have shown the influence of acoustic stimulation on visual responses across different species (Iurilli et al., 2012; Ibrahim et al., 2016; Chanauria et al., 2019; Bimbard et al., 2023; Williams et al., 2023). Investigations have also shown visual neurons exhibit plastic modifications in relation to auditory inputs (Iurilli et al., 2012; Ibrahim et al., 2016; Chanauria et al., 2019; Williams et al., 2023). Since there are cross-modal interaction between visual and auditory systems, here, we specifically hypothesized that the mechanisms of visual *vs.* sound adaptation (a protocol that induces plasticity in neurons well into adulthood) might be different in visual neurons. To this goal, in this investigation, we tested the effect of visual *vs.* sound adaptation across the supra- and infra-granular layers of the cat visual cortex by computing the AUC on a trial-by-trial basis [see section, Data analysis: area under the curve (AUC) computation]. We tested 45 and 48 neurons in the supra- and infra-granular layers, respectively, for the visual adaptation and 78 and 80 neurons, respectively for the sound adaptation.

### 4.1 Comparison between visual vs. sound adaptation: increased variance for sound adaptation

Our previous investigations have demonstrated that visual adaptation can lead to either attractive or repulsive shifts in visual neurons, whereas sound adaptation may change the selectivity of visual neurons to either side of the original preferred orientation. Examples of such shifts are shown in the **Supplementary** Figure 2. Here, we specifically thought that visual and sound adaptation may have distinct effects on the dendritic spines to potentiate the tuning shifts. Therefore, here we began by computing the area of the raw tuning curves for each of the trials pre-and post-adaptation. Figure 2 illustrates the results of this analysis. An example of the AUC computation for a trial is shown in the Supplementary Fig. 1.

**Figure 2:**
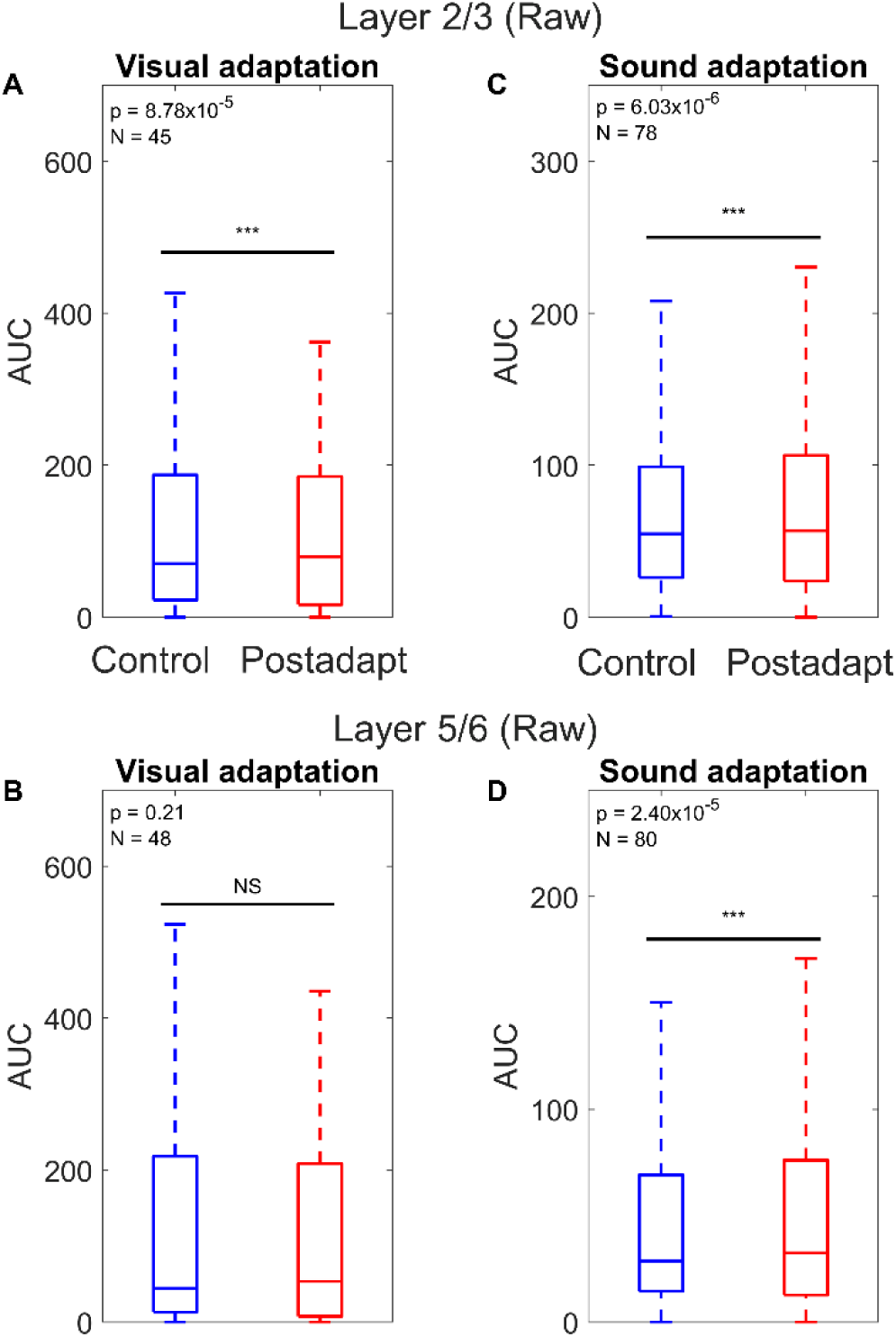
Boxplots presenting AUC of the raw tuned data observed in layers 2/3 and 5/6 neurons for visual vs. sound adaptation. **(A)** Visual adaptation: Significant difference in AUC for control (blue) vs. adaptation (red) for layer 2/3 neurons (N = 45; Wilcoxon signed-rank test, p = 8.78 x 10^-5^). **(B)** Visual adaptation: no significant difference in AUC for control vs. adaptation for the layer 5/6 (N = 48; Wilcoxon signed-rank test, p = 0.21). (**C)** Sound adaptation: significant difference in AUC for control vs. adaptation for layer 2/3 neurons (N = 78; Wilcoxon signed-rank test, p = 6.03 x 10^-6^). **(D)** Sound adaptation: significant difference in AUC for control vs. adaptation for layer 5/6 neurons (N = 80; Wilcoxon signed-rank test, p = 2.40 x 10^-5^).

For visual adaptation in layers 2/3 (Fig. 2A), we noticed a significant difference for AUC (N = 45, Wilcoxon signed-rank, p = 8.78x10^-5^) between pre- (mean = 136.80 ± 5.29, median = 70.50) and post-adaptation (mean = 143.42 ± 5.59, median = 79.50), whereas for the layer 5/6 (Fig. 2B) no significance (N = 48, control: mean = 135.59 ± 5.82, median = 44.25; post-adaptation: mean = 131.70 ± 5.67, median = 53.50; Wilcoxon signed-rank, p = 0.21) was observed.

Interestingly, in the case of sound adaptation a significant difference was observed across both layers: Layer 2/3 (Fig. 2C, N = 78, control : mean = 86.15 ± 2.14, median = 54.75; post-adaptation : mean = 92.27 ± 2.32, median = 56.70; Wilcoxon signed-rank, p = 6.03x10^-6)^ and Layer 5/6 (Fig. 2D, N = 80, control : mean = 61.18 ± 1.88, median = 28.69; post-adaptation : mean = 69.26 ± 2.45, median = 39.18; Wilcoxon signed-rank, p = 2.40x10^-5^). Notably, for visual adaptation there was no significant difference in difference in variance across both layers (Layer 2/3, Two-sample F-test, p = 0.06; Layer 5/6, Two-sample F-test, p = 0.37), whereas for sound adaptation, a significant increase was noticed post-adaptation across both layers (Layer 2/3, Two-sample F-test, p = 2.35x10^-4^; Layer 5/6, Two-sample F-test, p = 3.24x10^-9^). This difference in variance implies that sound leads to more perturbations on the dendritic synapses, thereby leading to more differences in the firing of neurons from one trial to another.

We then hypothesised that adaptation may impinge specifically upon the original optimal (control) and new optimal (after adaptation) orientation, since most of the post-adaptation effects are restricted to the original and the newly acquired orientation (Ghisovan et al., 2009; Bharmauria et al., 2022).

Therefore, we proceeded by centering the responses of neurons on the original and novel optimal orientations (computed as mean of responses across 25 trials) of neurons in the control and post-adaptation conditions, respectively. Again, we obtained similar results as for the uncentered raw data (Fig. 2). In the case of visual adaptation, a significant difference was noticed in layer 2/3 (Fig. 3A, N = 45, control: mean = 157.44 ± 6.04, median = 82,00; post-adaptation: mean = 163.91 ± 6.39, median = 89.00; Wilcoxon signed-rank test, p = 2.17x10^-^) for AUC between the pre- and post-adaptation conditions, whereas for the layer 5/6 no significance (Fig. 3B, N = 48, control : mean = 155.89 ± 6.70, median = 48.00; post-adaptation : mean = 150.98 ± 6.53, median = 57,00; Wilcoxon signed-rank test, p = 0.45) was observed. Whereas in the case of sound adaptation, a significant difference was observed in both layers post-adaptation: Layer 2/3 (Fig. 3C, N = 78, control: mean = 97.81 ± 2.42, median = 62.74; post-adaptation: mean = 105.53 ± 2.62, median = 67.75; Wilcoxon signed-rank, p = 7.27x10^-6^) and Layer 5/6 (**Fig 3D**, N = 80, control: mean = 71.28 ± 2.19, median = 32.96; post-adaptation : mean = 79.42 ± 2.45, median = 39.18; Wilcoxon signed-rank test, p =1.72x10^-4^). Again, there was no significant difference in variance for visual adaptation (Layer 2/3, Two-sample F-test, p = 0.06; Layer 5/6, Two-sample F-test, p = 0.38) but for sound adaptation a significant difference was noticed (Layer 2/3, Two-sample F-test, p = 7.03x10^-4^; Layer 5/6, Two-sample F-test, p = 2.63x10^-7^). Collectively, the above analysis (Fig. 2**-3)** suggests divergent effects of either protocol on V1 neurons.

**Figure 3:**
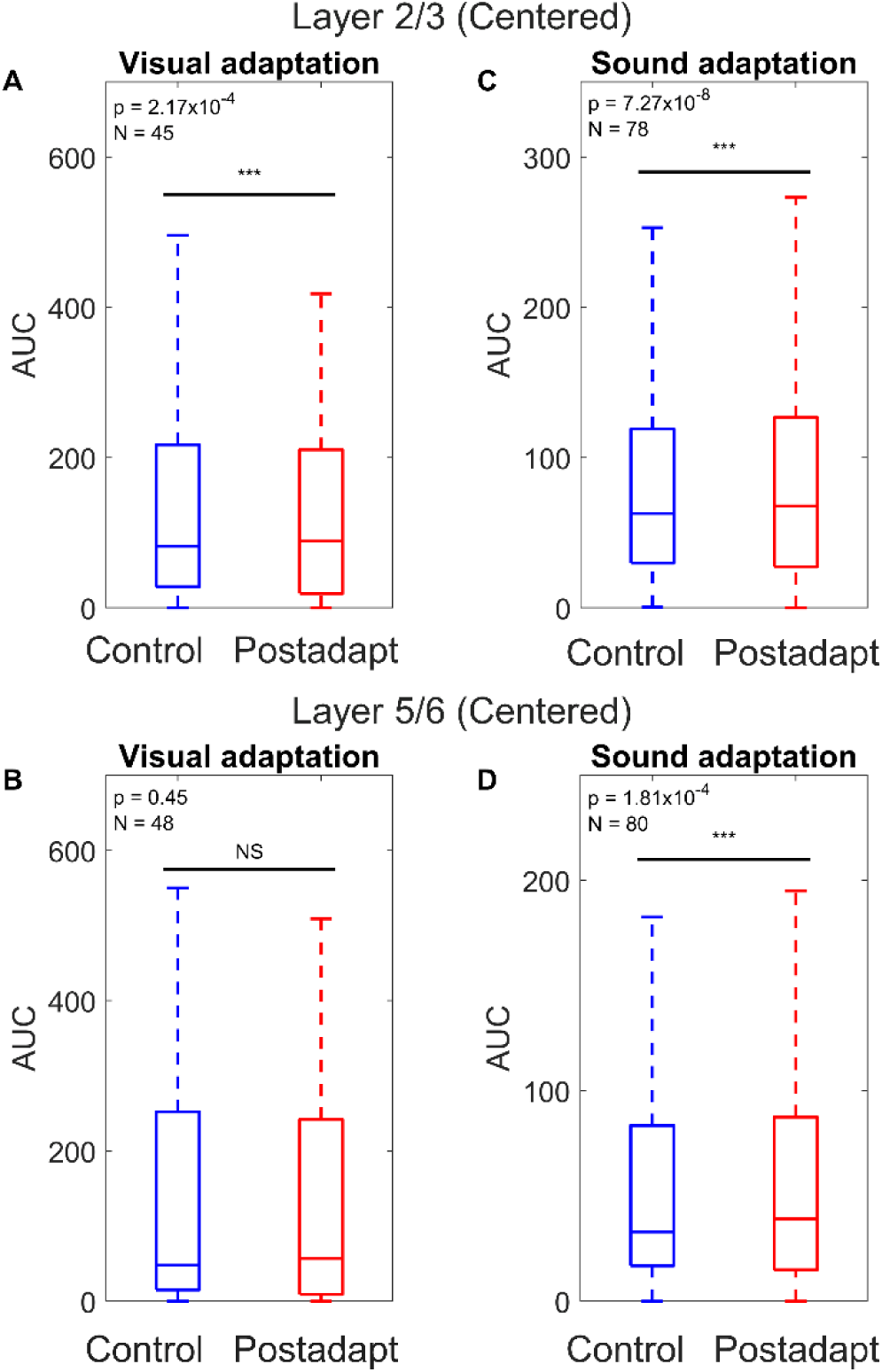
Boxplots presenting AUC of the cantered tuning curves in layers 2/3 and 5/6 for visual vs. sound adaptation. **(A)** Visual adaptation: Significant difference in AUC for control (blue) vs. adaptation (red) for layer 2/3 neurons (N = 45; Wilcoxon signed-rank test, p = 2.17x10^-4^). **(B)** Visual adaptation: no significant difference in AUC for control vs. adaptation for the layer 5/6 (N = 48; Wilcoxon signed-rank test, p = 0.45). (**C)** Sound adaptation: significant difference in AUC for control vs. adaptation for layer 2/3 neurons (N = 78; Wilcoxon signed-rank test, p = 7.27x10^-6^). **(D)** Sound adaptation: significant difference in AUC for control vs. adaptation for layer 5/6 neurons (N = 80; Wilcoxon signed-rank test, p = 1.72x10^-4^).

Finally, in line with above predictions, to test the modulations around the original and novel orientations, we plotted the firing rate as a function of orientations (centered data) across all the trials for all the neurons in both conditions. Figure 4 demonstrates these results. For visual adaptation, in layer 2/3 (Fig. 4A), there was no significant difference between the control and post-adaptation conditions (N = 45; Kolmogorov–Smirnov test, p = 0.13). Similarly, for layer 5/6 (Fig. 4B), no significant difference was observed (N = 48; Kolmogorov–Smirnov test, p = 0.39). Notice the overlap of the control (blue) and post-adaptation (red) curves in both cases, indicating that the effect of visual adaptation was rather restricted to the original and novel orientations (Bharmauria et al., 2022).

**Figure 4:**
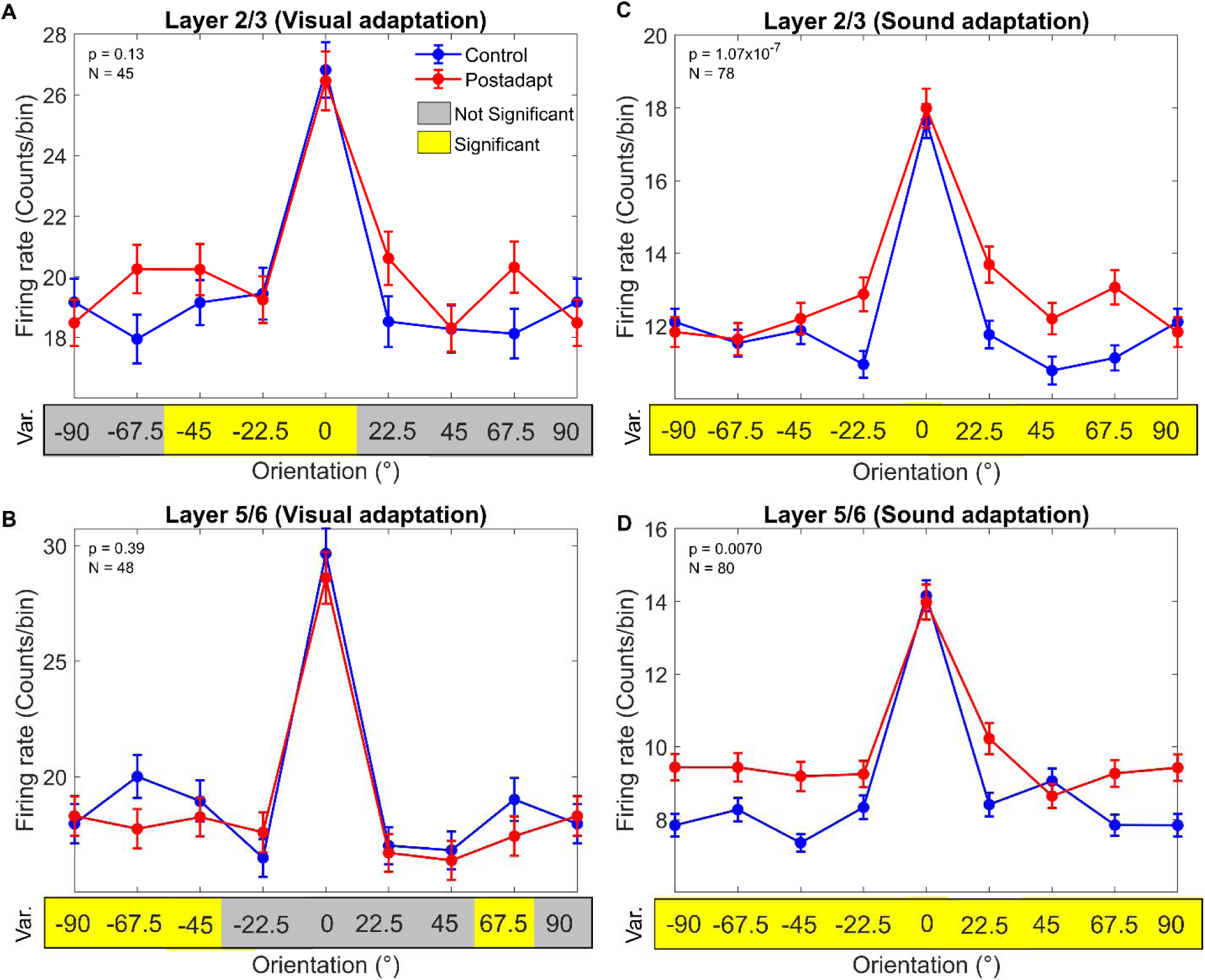
Comparison between centered tuning curves for visual vs. sound adaptation. **(A-B)** In the visual condition, the control (clue) and post-adaptation (red) data overlap and show no significant difference in both layers (Layer 2/3, Kolmogorov–Smirnov test, p = 0.13; Layer 5/6 Kolmogorov–Smirnov test, p = 0.39). The yellow / grey squares on the x-axis depict the significance / non-significance (Two-sample F-test, p < 0.05) for variance at that orientation. Most of the orientations exhibit equal amounts of variance (grey squares) pre- and post-adaptation. **(C-D)** In the sound condition, in both layers, neurons showed significantly broader tuning (Layer 2/3, Kolmogorov–Smirnov test, p = 1.07x10^-7^; Layer 5/6 Kolmogorov–Smirnov test, p = 0.007) accompanied with higher firing rates at the flanking orientations after sound adaptation. All the orientations are characterized by an increase of variance post-adaptation, as depicted by yellow squares. The data are presented as mean ± sem.

Remarkably, when we performed the same analysis for sound adaptation, a significant difference was observed in both layers: Layer 2/3 (Fig. 4C, N = 78; Kolmogorov– Smirnov test, p = 1.07x10^-7^) and Layer 5/6 (Fig. 4C, N = 80; Kolmogorov–Smirnov test, p = 0.007). Here, there was a pronounced broadening of the tuning curve post-adaptation (red) compared with the control (blue) condition, with visible response gain in the flanks across both layers, suggesting an increased synaptic drive at the flanks (Jia et al., 2010; Wilson et al., 2016; Bharmauria et al., 2019).

These results are further corroborated by our analysis on the variance on an orientation-by-orientation basis for both conditions. The significant difference (Two-sample F-test, p < 0.05) is indicated by the yellow and grey squares (for better visual comparison) along the x axis for each orientation: yellow colour represents significant difference in variance for that orientation and grey depicts a non-significant difference. Note that in the case of visual adaptation (Fig. 4A**-B**, see the x axis), most of the squares are grey, implying that at most of the orientations, there is almost equal synaptic drive pre- and post-adaptation. Contrastingly, all the squares are yellow at all the orientations for the sound condition (Fig. 4C**-D**), suggesting an increased synaptic drive from the control to post-adaptation condition. All the p-values for the respective orientations (for variance and distribution difference) are shown in the **Supplementary** Figure 3.

Since neurons may fire differently from one trial to another, we also performed the same analysis (**Supplementary** Fig. 4) by centring the tuning curves on a trial-by-trial basis. Again, we obtained the same results, as above: no significant difference was observed for visual adaptation across both layers, whereas sound adaptation significantly broadened the tuning curves. In conclusion, these results strongly indicate that sound adaptation affects the underlying neural dendritic structure leading to the broadening of raw tuning curves (increased bandwidth), thus the overall increase in their areas, i.e., variance from the control to post-adaptation condition.

## 4. Discussion

The findings presented in the study shed light on the intricate relationship between visual and auditory stimuli and their impact on the neural processing within the cat visual cortex. Specifically, we report that sound adaptation led to the broadening of the novel raw tuning curves, characterized by increased variance. Based upon previous reports (Jia et al., 2010; Wilson et al., 2016), we speculate that sound modifies the input-output function of neuronal dendrites much more non-linearly than the visual adaptation.

### 4.1 Comparative Results: visual *vs.* sound adaptation

First, these experiments were done in anaesthetized cats, so all the fluctuations related to attentional parameters are ruled out (Gray and Singer, 1989; Bharmauria et al., 2015). Notably, while visual adaptation led to both attractive and repulsive shifts in visual neurons (Ghisovan et al., 2009; Bachatene et al., 2015c; Chanauria et al., 2016), sound adaptation tuning shifts towards different orientations (Chanauria et al., 2019). One of the key observations from the study is the differential impact of adaptation across cortical layers. In the case of visual adaptation, although significant differences were predominantly observed in layer 2/3 for AUC, but for layer 5/6, no difference was noticed. Whereas for sound adaptation changes were significantly elicited across both layers.

In the sensory cortex, layer 2/3 receives direct vertical inputs from layer 4 (that receives information from the later geniculate nucleus), and layer 2/3 relays these inputs to the layer 5/6 (Gilbert and Wiesel, 1989; Hirsch and Martinez, 2006; Bachatene et al., 2012a). Our recordings were performed using linear probes in the cortex (possibly recording an orientation column); therefore, we expect to record similar behavior between layers 2/3 and 5/6 neurons pre-and post-adaptation. Consistent with this, our results on visual adaptation suggest that, indeed, adaptation aftereffects in layers 2/3 and 5/6 go hand-in-hand, as revealed previously (Chanauria et al., 2016). Specifically, similar proportion of attractive and repulsive shifts in both layers across different cell types (Chanauria et al., 2016). On the contrary, after sound adaptation (Chanauria et al., 2019) in layer 2/3, neurons exhibited a notable inclination to adjust their preferred orientation towards horizontal axis, whereas cells in the layer 5/6 displayed a preference shift spanning the entire orientation spectrum. Studies in cats have demonstrated that orientation selectivity begins at the cortical level rather than at the LGN level (Dragoi et al., 2001; Bouchard et al., 2008), suggesting that these disparate tuning changes to be of cortical origin. Furthermore, using OSI computation, cellular bandwidths increased with sound for only layer 5/6, whereas in layer 2/3, an opposite pattern was observed (Chanauria et al., 2016). This allowed us to argue that both layers work independently of each other, in response to sound, irrespective of the position (lateral vs. frontal) of the sound presentation.

In spite of the differences documented between layers 2/3 and 5/6 to sound adaptation (Chanauria et al., 2019), here, with AUC calculations, we notice that both layers exhibited significant broadening of their raw tuning curves accompanied with increased variance at all orientations. Auditory inputs arrive at layers 1 and 2/3 of the V1 from the primary auditory cortex (A1) (Ibrahim et al., 2016; Deneux et al., 2019; Garner and Keller, 2022). These direct inputs to the layer 2/3 may explain the bias toward the horizontal axis in our case but not in the layer 5/6. Given that layer 5/6 receives input from layer 2/3, how do the layer 5/6 neurons display a different post-adaptation behaviour than layer 2/3 neurons, although both are characterized by similar variance pattern around the optimal orientation (Fig. 4C**-D**)? We discuss that in the following.

### 4.2 Neural mechanism: synaptic & excitation/inhibition changes across layers

Figure 5 demonstrates a model of these distinct tuning changes with respect to the visual and sound adaptation. In the case of visual adaptation (Fig. 5A), layer 2/3 neurons exhibit tuning changes from a large range of orientation (light blue) to a large range of orientation (light red) tuning curves; notably, without any change in the average tuning curves (dark blue) and (dark red) following adaptation. This is characterized by an increased inhibition (blue and arrows)on the flanks pre-and post-adaptation from the nearby neurons and lateral inhibition from the other columns (Wei et al., 2022), thereby sharpening the tuning curve of the neuron. In fact, studies have shown that the fundamental organization of orientation maps is preserved pre- and post-adaptation despite changes in the orientation map (Dragoi et al., 2000; Cattan et al., 2014; Bachatene et al., 2015a; Li et al., 2017). Importantly, although visual adaptation changes the responses of retinal ganglion cells (RGC) and the lateral geniculate nucleus (LGN), it does so without changing the contrast gain control of V1 than the RGCs and the LGN (Solomon et al., 2004; Camp et al., 2009; Dhruv and Carandini, 2014; Li et al., 2017), thus maintaining the population homeostasis after plastic modifications in the V1 (Benucci et al., 2013; Bachatene et al., 2015b).

**Figure 5:**
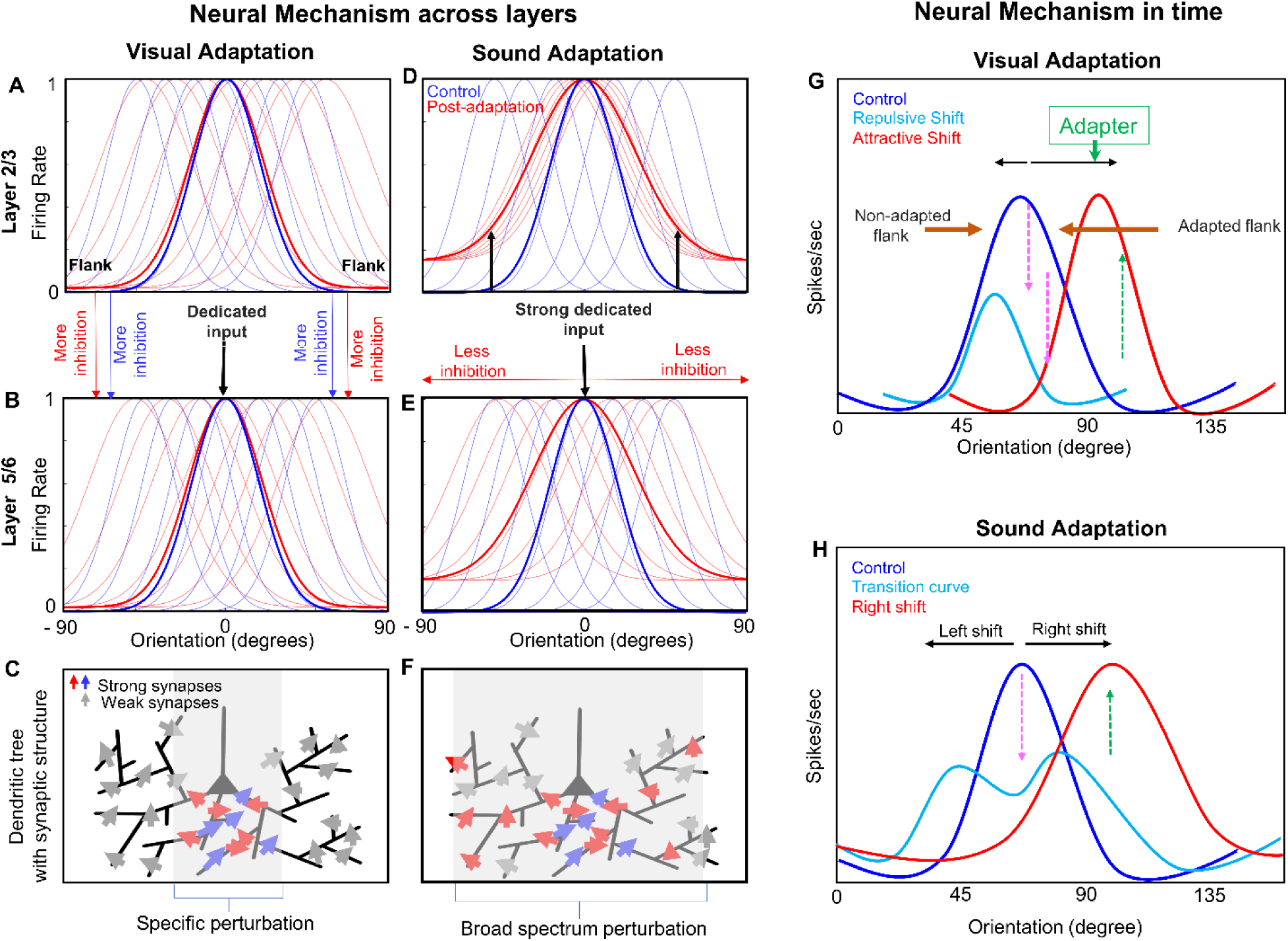
Comparable mechanisms of the acquisition of novel tuning curves after visual and sound adaptation. **(A)** Similar range of layer 2/3 tuning curves pre- (blue) and post-adaptation (red) in relation to the visual input. The thick curves represent average tuning curves pre- and post-adaptation. Note that their average tuning is similar. **(B**) A dedicated oriented synaptic input (pre- and post-adaptation, indicated by red and blue arrows in **C**) arrives from the optimal orientation of layer 2/3 at layer 5/6, thereby changing the tuning of layer 5/6 neurons through specific dendritic perturbations, characterized by more inhibition on the flanks in both conditions. Neurons change their tuning to the either side in relation to adaptation. **(D)** Conversely, during sound adaptation, the input arrives from the primary auditory cortex (A1), thus changing the tuning of the V1 neurons toward the horizontal axis (note all neurons tuned to 0 °). This strong input fatigues the synapses corresponding to the optimal orientation, concurrently with the release of inhibition on flanks. **(E)** The strong 0 ° input from the layer 2/3 to layer 5/6, cascades into suppression of responses at the optimal orientation with the release of inhibition on flanks (with a broad spectrum of dendritic perturbation as in **F**). This leads to the broadening of tuning curves of layer 5/6. Notably, cross-orientation inhibition may subserve the emergence of tuning and shifts in either direction. **(G)** Push-pull mechanism of orientation tuning shifts during visual adaptation. First, decreased responses (pink arrows) at the adapted flank of the original tuning curve (blue), caused by the adapter (downward green arrow), lead to a repulsive shift as synapses exhaust. This shift results in the dominance of the opposing flank. Over time, selective response increase (green upward arrow) near the adapter leads to an attractive shift, with concurrent suppression of neuronal activity at the non-adapted flank. **(H)** Similarly, during sound adaptation, continuous drive from the A1 at layer 2/3 may suppress responses at the optimal tuning curve, causing a brief sliding of the tuning curve in either direction. Eventually, dominance of synaptic drive toward one side leads to the emergence of a novel tuning curve. Shifts in layer 2/3 tend toward the horizontal axis, while in layer 5/6, equal left- and right-ward shifts may occur.

During visual stimulation pre- and post-adaptation, when the information from the layer 2/3 is relayed to the layer 5/6 neurons (Fig. 5B), it begins with a dedicated optimal drive (downward black arrow) to the pyramidal neurons, attributed to only a select group of strong synapses representing particularly oriented inputs (blue and red arrows, **Fig 5C**). The synapses corresponding to other oriented inputs are much weakly driven (grey arrows). As pointed above, this features strong inhibitory input acting upon them, thereby allowing the selection of the optimal stimulus (Harth and Pertile, 1972; Scholl et al., 2013; Wei et al., 2022). Briefly, the inhibition/excitation balance is not disturbed from pre- to post-adaptation in the visual condition, within and between the supra- and infra-granular layers. This allows for a swift transfer to the adaptive code, highlighted by changes mostly restricted to the original and novel orientations, devoid of much disturbance on the flanks — explaining less variance.

Conversely, in the sound condition, there is a new dynamic that is achieved following adaptation. In our experiment, after the layer 2/3 neurons receive the input form the A1, somehow the cyclical nature of our sound adapter alters the tuning of curves toward the horizontal axis, i.e., toward 0° orientation (Fig. 5D). Since most of the neurons now have achieved a 0°orientation, there is a hyperdrive from the layer 2/3 acting upon entire hyper-column(s) of the layer 5/6. This strong dedicated input has implications that are twofold: 1) the repetitive incoming drive from layer 2/3 may suppress the response of the layer 5/6, thus leading to the release of inhibition on the flanks. 2) The weakened inhibition on the flanks in conjunction with increased cross inhibition between the columns may lead to emergence of the new orientation tuning curves with a broad spectrum **(Fig 5E).** Overall, there is much more activation of synapses corresponding to other orientations in the case of sound adaptation (Fig. 5F). There is a possibility that this form adaptation occurs without the involvement of RGCs and the LGN (Solomon et al., 2004; Camp et al., 2009; Dhruv and Carandini, 2014; Li et al., 2017), thus substantially altering the population homeostasis (Benucci et al., 2013; Bachatene et al., 2015b). The feedforward and feedback loops between the layers may also be involved in the refinement of these novel features (Yoshimura et al., 2000; Markov et al., 2014).

### 4.3 Neural mechanism in time

To explain our visual adaptation results, we put forth a push-pull mechanism for orientation selection (Ghisovan et al., 2009; Bharmauria et al., 2019, 2022), consisting of two independent but related mechanisms. First, the decline of responses (downward dashed pink arrows) at the adapted flank of the original tuning curve (blue) (Fig. 5G) due to the adapter (green downward arrow) results in a repulsive shift (cyan) as the tuning curve slides away from the adapter because of the exhaustion of synapses (Carandini and Ferster, 1997; Thomson and Deuchars, 1997; Harris et al., 2000; Chelaru and Dragoi, 2008). This results in the opposing flank being dominant. With time, conversely, a selective increase in responses (upward green arrow) near the adapter results in an attractive shift (Fig. 5H, red curve) from a subsequent suppression of neuronal activity at the non-adapted flank. Notably, the establishment of novel tuning curves may be significantly influenced by lateral contacts (mutual inhibition / excitation of cells) between orientation columns as noted above (Carandini et al., 1998; Crook et al., 1998; Wei et al., 2022).

Within the same framework, we explain our results on sound adaptation: first, the continuous drive arriving from the A1 at layer 2/3 (Ibrahim et al., 2016; Deneux et al., 2019; Garner and Keller, 2022) may lead to a suppression of response at the optimal tuning curve, thus causing a brief sliding of the tuning curve in either direction (cyan curve) of the optimal tuning. With time, as the synaptic drive to one side of the curve dominates over the other flank, this would lead to the emergence of a novel tuning curve toward the dominant side of the curve: in this case, to the right (red curve). Notably, in the layer 2/3 most of the shifts may be potentiated toward the horizontal axis but as the input is relayed to the layer 5/6, an equivalent drive (with factors as mentioned in the above section) may engender left- and right-ward shifts in equal proportions.

Importantly, distinct cortical regions, layers, and cell subtypes may adapt differently from one another due to differing dendritic topologies (Ahmed et al., 1998; La Camera et al., 2006), thus their roles across layers need to be further investigated. For example, compared with layer 6 pyramidal neurons, layer 3 pyramidal neurons adapt far more quickly (Ahmed et al., 1998). Neurons in the cat’s layer 2/3 of V2 are more impacted by adaptation than layer 5/6 (Lussiez et al., 2021). Pyramidal cells (regular spiking) adapt significantly more quickly than putative interneurons (rapid spiking) (La Camera et al., 2006; Mensi et al., 2012), and that the magnitude of shifts in pyramidal cells is greater than that of interneurons (Bachatene et al., 2012b). Significantly, after adaptation, feedback from higher visual areas may control the learning of new tunings even more (Galuske et al., 2002; Keller et al., 2020; Huang et al., 2022).

### 4.4 Functional benefit to the cross-modal systems(s)

Even before a child is born, sound is available to the brain, thus it plays a substantial role into the adulthood for cross-modal interactions and abstract representation of things to the brain and help one visualize them (Movalled et al., 2023). In our case, the broadening of the V1 tuning curves during sound adaptation may possibly be directly associated with the nature of sound stimulus presented, its spectral or temporal features. Since visual neurons needed to encode different frequencies of the sound during sound adaptation, increased bandwidth in cortical layers may allow effective translation of multifrequency stimulus to the visual system (Knöpfel et al., 2019). Moreover, this broad tuning enables neurons to enhance their ability to encode or bind complex information during sound adaptation (Pearson, 2019; Opoku-Baah et al., 2021). In fact, when the auditory perception / imagery information is available to the brain, the early visual cortex receives non-retinal information, thereby, enabling the representation of common abstract information (Vetter et al., 2014). Another possibility is that sound was presented in the overlapping receptive fields of the auditory cortex that evoked distinct neuronal responses in the layer 2/3 of the V1, recorded as broad of tuning curves unlike sharper ones recorded during visual adaptation. Additionally, a sound stimulus presented in the overlapping receptive fields may trigger an ambiguous, yet significant sound-induced visual response, that perhaps could only be represented with the broadening of V1 tuning curves. How else could a visual neuron encode dynamic features of the sound stimulus?

It was shown in mice that presentation of sound burst drove a suppressive effect in V1 (Iurilli et al., 2012), driven by corticocortical excitatory projections from the A1 to supra-granular neurons. Therefore, along with the characteristics of the sound, the duration of its presentation has an equal role to play. Furthermore, the rapid and delayed mechanisms of sound representation in the visual cortex may enable us to adapt to different stimuli and environments.

## 5. Conclusion

In summary, this study enhances our comprehension of sensory adaptation by illustrating the unique impacts of visual and auditory stimuli on neurons in the cat visual cortex. Through elucidating the varied mechanisms governing sensory plasticity, these findings propel our understanding of cortical processing forward and set the stage for future explorations into the integration of multiple senses and their influence on behavior. The study also necessitates meticulous controls to differentiate between auditory signals and neural activity associated with internal states and behaviors. Further studies need to be done to understand the contribution of motor movements to neural responses evoked by sound (Ibrahim et al., 2016; Bimbard et al., 2023; Williams et al., 2023). Overall, this investigation underscores the importance of comprehensive inquiries into the mechanisms governing sound-induced activity within the visual cortex, considering the diverse array of factors influencing neural reactions. Ultimately, these findings offer foundational insights into the distinct mechanisms of plasticity within the sensory cortex, potentially ensuring its readiness across various abstract representations corresponding to visual stimuli.

## Contributions

YYB analysed the data, made the figures, and contributed to writing. SPLH helped in data analysis. NC performed the experiments and contributed to writing. OF contributed to data analysis, editing and writing. SM contributed to writing and editing of the manuscript. VB wrote the first draft, contributed to data analysis, editing, figures and supervised the study.

## Acknowledgements

This study was supported by a Natural Sciences and Engineering Research Council of Canada (NSERC) grant to SM.

