## Supplementary data for "Distinct mechanisms of visual and sound adaptation in the cat visual cortex"

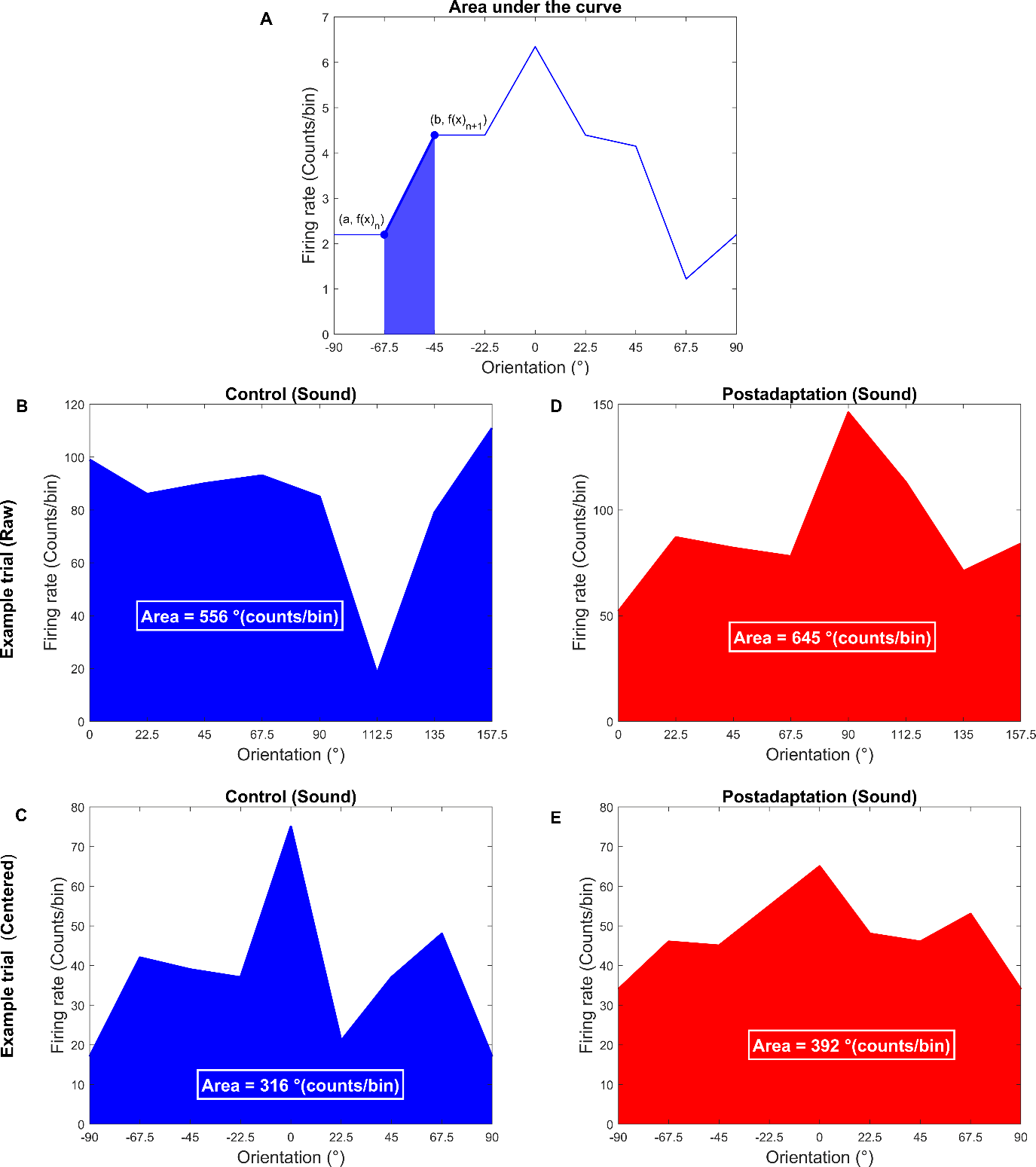

***Supplementary Fig. 1****: Area under the curve (AUC) computation.* ***(A)*** *Computation of AUC (see methods).* ***(B-C)*** *An example of AUC pre-* ***(B)*** *and post-adaptation* ***(C)*** *for a trial (uncentered data) in the sound condition.* ***(D-E)*** *An example of AUC pre-* ***(B)*** *and post-adaptation* ***(C)*** *for another trial (centred data) in the sound condition.*

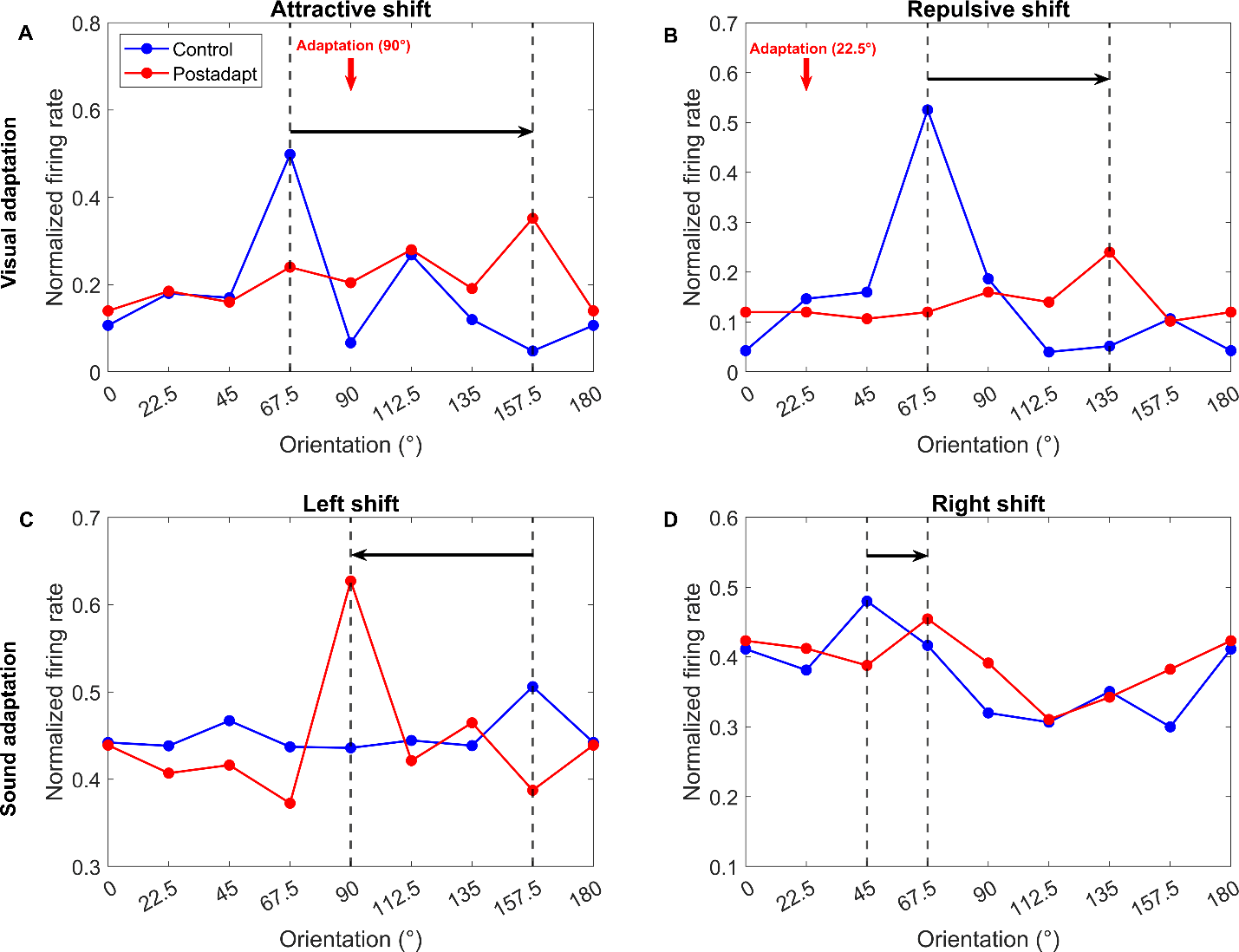

***Supplementary Fig. 2****: Examples of tuning shifts (raw curves) for visual vs. sound adaptation.* ***(A)*** *An example of an attractive shift for a visual neuron in the cat V1. Blue line stands for the control and red line represents post-adaptation tuning. From control (67.5 °) to post-adaptation (157.5 °), the neuron displayed an attractive shift of 90 ° in orientation tuning, i.e., in the direction of the adapter (indicated by red downward arrow).* ***(B)*** *An example of a repulsive shift. Here the neuron displayed a shift (67.5 °) in the direction opposite to the adapter.* ***(C)*** *An example of a leftward shift (67.5 °) for sound adaptation.* ***(D)*** *An example of a rightward shift (22.5°) for sound adaptation.*

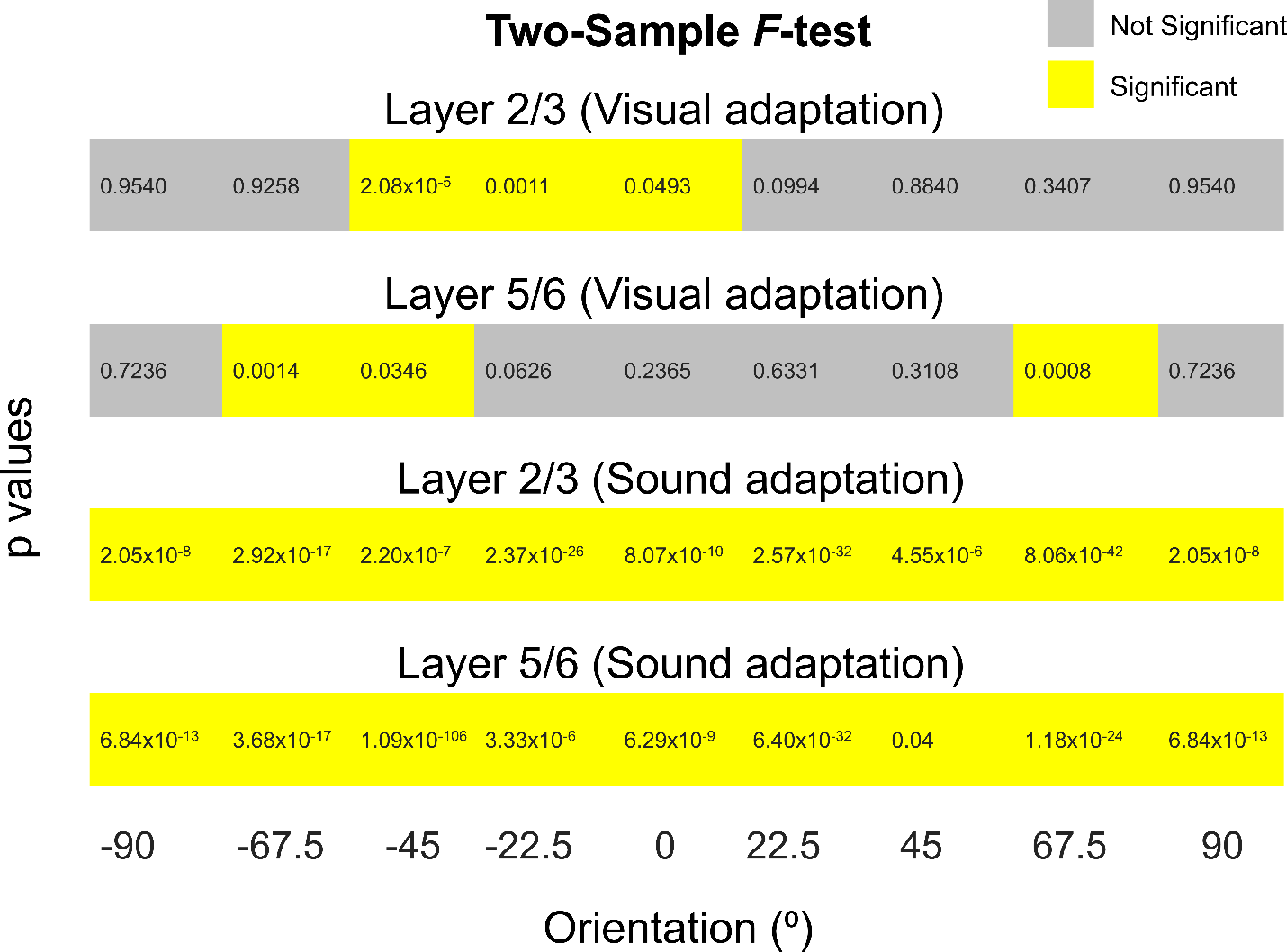

***Supplementary Fig. 3****: p- values (Two-sample F test for variance) for comparison between control and post-adaptation for visual and sound and adaptation. Yellow and grey represent significance and non-significance, respectively.*

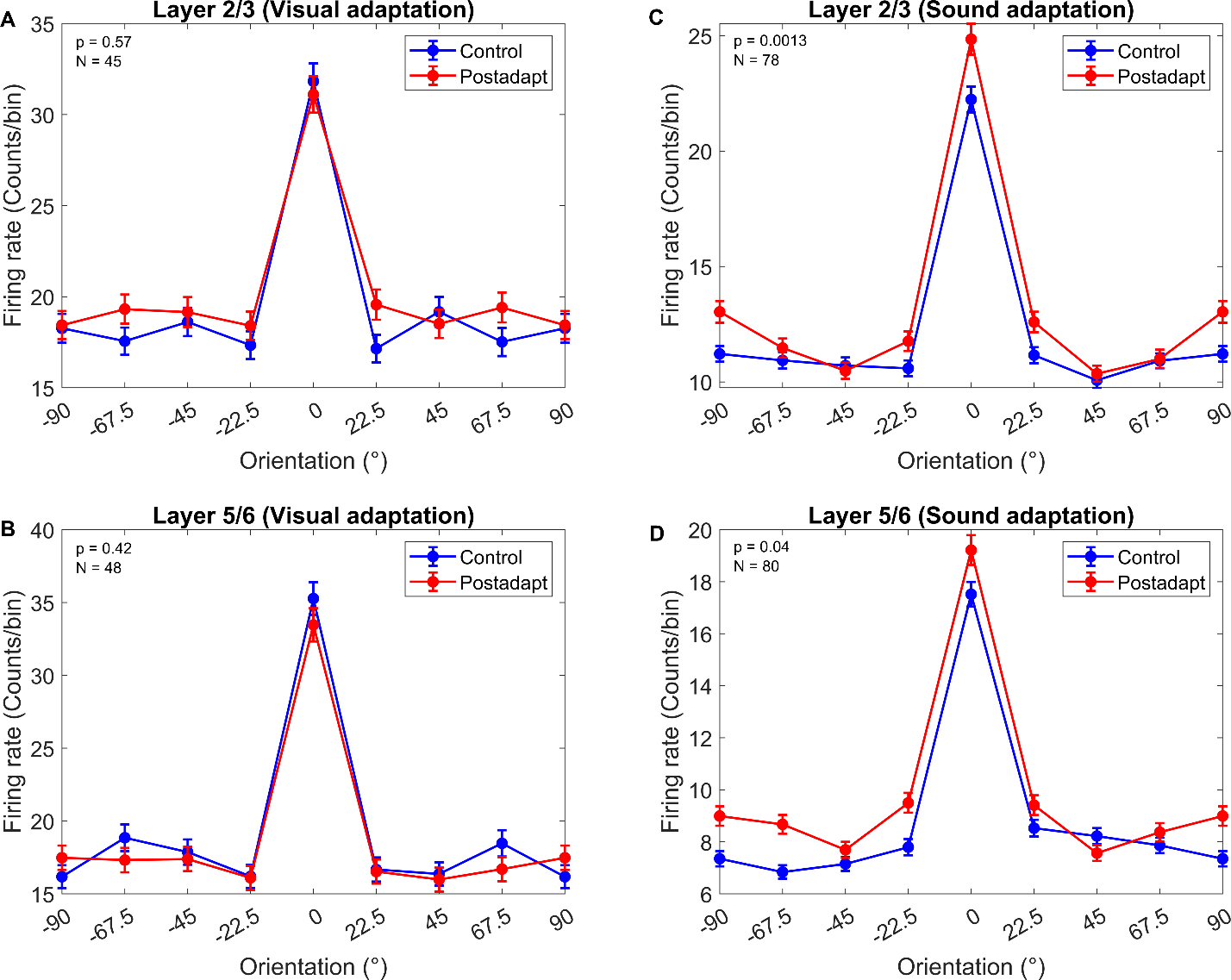

***Supplementary Fig. 4****: Comparison between centered tuning curves for visual vs. sound adaptation (on a trial-by-trial basis)* ***(A-B)*** *In the visual condition, the control (clue) and post-adaptation (red) data overlap and show no significant difference in both layers (****C****-****D).*** *In the sound condition, in both layers, neurons showed significantly broader bandwidths accompanied with higher firing rates at the flanking orientations after sound adaptation. The data are presented as mean ± sem.*
